# Bacteria are required for regeneration of the *Xenopus* tadpole tail

**DOI:** 10.1101/319939

**Authors:** Thomas F. Bishop, Caroline W. Beck

## Abstract

The impressive regenerative capabilities of amphibians have been studied for over a century. Although we have learnt a great deal about regenerative processes, the factors responsible for the initiation of regeneration have remained elusive. A previous study implicated reactive oxygen species (ROS) and the ROS-generator, NADPH oxidase (Nox), in *Xenopus* tadpole tail regeneration. In this study we suggest that Nox is expressed as a consequence of NF-*κ*B transcription factor activity and that ROS produced by Nox, in turn, help to maintain the activity of NF-*κ*B, forming a positive-feedback loop. Microorganisms were found to be required for regeneration through binding to toll-like receptors (TLR). NF-*κ*B is a downstream component of TLR pathways and its activation through TLR stimulation could jump-start the positive-feedback loop. These findings provide potential targets for the activation of regeneration in non-regenerative animals.

## Introduction

Amphibians have remarkable regenerative capabilities, but the mechanisms they use to initiate and maintain regeneration are still largely unknown. It has been previously shown by Love et al (2013) that reactive oxygen species (ROS) are continually produced and required for successful tadpole tail regeneration ***Love et al.*** (***2013***). Production of ROS and tadpole tail regeneration are prevented by NADPH oxidase (Nox) inhibitors, suggesting Nox complexes as the source of ROS ***Love et al.*** (***2013***). However, the role of ROS and the mechanism of their sustained production throughout regeneration were uncertain.

NF-*κ*B is a rapid-acting transcription factor with the potential to dramatically alter the activity and function of a cell ***Sun and Andersson*** (***2002***). NF-*κ*B is necessary for maintaining the undifferentiated state of human embryonic stem cells ***Deng et al.*** (***2016***), human induced pluripotent stem cells ***Takase et al.*** (***2013***) and mesenchymal stem cells ***Chang et al.*** (***2013***), suggesting it could be involved in maintaining the de-differentiated state of regeneration blastema cells. In the absence of an activating signal, NF-*κ*B is sequestered in the cytoplasm by I*κ*B (inhibitor of NF-*κ*B), preventing its nuclear localisation and activity. The I*κ*B kinase (IKK) complex inhibits I*κ*B in response to multiple extracellular stimuli, but ROS can also inhibit I*κ*B ***Reuter et al.*** (***2010***). Nuclear NF-*κ*B directly activates transcription of several genes encoding Nox proteins ***Manea et al.*** (***2010***); ***Morgan and Liu*** (***2011***), so could thereby facilitate ROS production.

Here, we suggest that a positive-feedback loop involving NF-*κ*B could maintain ROS levels in regenerating amphibian appendages. Intracellular ROS can inhibit I*κ*B, which would result in increased NF-*κ*B activity. Active NF-*κ*B could then facilitate the continual production of ROS by activating the transcription of Nox-encoding genes. Intriguingly, we also demonstrate that microorganisms can play a role in the initiation of tadpole tail regeneration. Microorganisms present on the skin of tadpoles offer sources of ligands for toll-like receptor (TLR) pathway activation and consequently, IKK complex activity. We suggest that this mechanism leads to NF-*κ*B activation at the wound site. Finally, we provide evidence that continual expression of the genes encoding Nox4 in limb bud blastema cells and Nox2 in professional phagocytes correlates with sustained NF-*κ*B activity.

## Results

### Nox activity is required and NF-*κ*B is activated within hours of amputation

It has been previously shown that ROS concentration greatly increases around the site of injury within 20 minutes after tadpole tail amputation, and is maintained at high levels throughout the process of regeneration ***Love et al.*** (***2013***). Raising tadpoles in the presence of the Nox inhibitor, DPI, for 3 days following tail amputation prevents regeneration ***Love et al.*** (***2013***). This shows that ROS are important for regeneration, but does not differentiate between the initial burst and the maintenance of ROS production. To see if ROS production was required during the initial hours following amputation, shorter DPI treatment durations were tested. Treatment with DPI beginning at 1 hour before amputation and terminating 1 hour afterwards had no effect on regeneration. However, tadpole tail regeneration was significantly reduced following DPI treatment for 3, 6 and 12 hours post-amputation (hpa) (Fig 1a). Longer treatments resulted in greater reductions in regeneration, with post-amputation treatments of 6 and 12 hours reducing regeneration scores to less than half of their respective controls. This confirms the previous finding that Nox activity is required for regeneration of the tail as early as 3 hours after tail amputation in *Xenopus* ***Love et al.*** (***2013***), and further identifies the early role of Nox in sustaining ROS levels.

**Figure 1.**
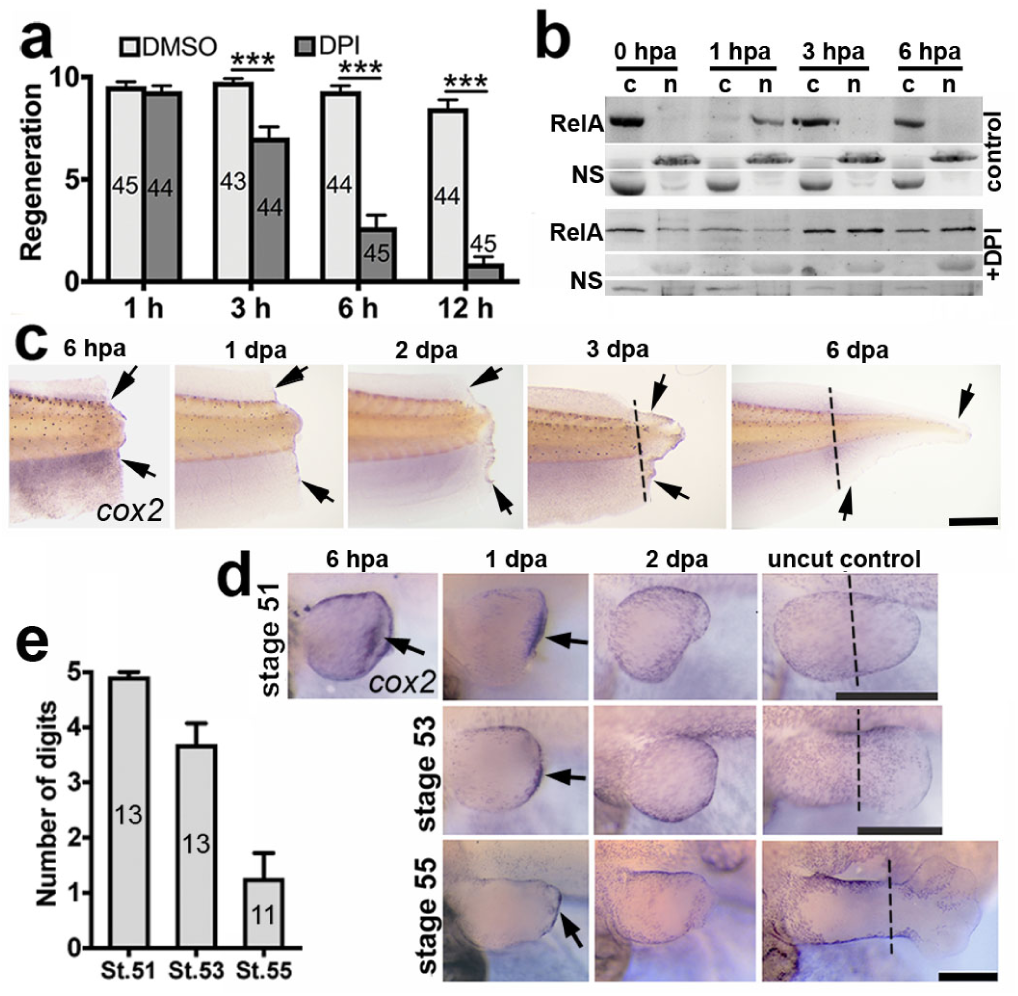
*Nox* and NF-*κ*B are active within hours after amputation. **a**, Mean tail regeneration scores + s.e.m of stage 43 tadpoles treated with DPI or DMSO control for 1 h before amputation and the indicated hpa. Sample numbers are indicated inside bars. *p* values are based on ordinal logistic regressions: *** *p* < 0.001. **b**, Western blots showing RelA in cytoplasmic C and nuclear N protein fractions from tail sections at indicated hpa ± DPI. Non-specific NS bands are included as loading controls. **c-d**, *In situ* hybridisations, using a *cox2* probe, of amputated tails (**c**) and hindlimbs (**d**). The stage at amputation and fixation times (hpa/dpa) are labelled. Black arrows indicate staining and dashed lines indicate the amputation planes in unamputated controls. Scale bars: 500 µm. **e**, Mean regeneration scores of hindlimbs amputated at the indicated stages + s.e.m. with sample numbers indicated inside bars. Data for graphs a and e can be found in supplemental file S1.

Inactive NF-*κ*B is sequestered in the cytoplasm and translocates to the nucleus upon activation. A Western blot of nuclear and cytoplasmic protein extracts from tadpole tail sections showed RelA (p65 subunit of NF-*κ*B) predominantly in the cytoplasm of uninjured tails and in the nucleus at 1 hpa (Fig 1b). By 3 hpa, it was once again located predominantly in the cytoplasm. The initial surge of ROS, where ROS were observed far beyond the amputation plane ***Love et al.*** (***2013***), could therefore account for the rapid NF-*κ*B translocation to the nucleus, suggesting that this key transcription factor is activated rapidly upon tail amputation. In the presence of DPI (Fig 1b), RelA was detected predominantly in the cytoplasmic fraction at 1 hpa, suggesting the involvement of ROS produced by Nox. The initial, widespread NF-*κ*B translocation may not be essential as DPI did not prevent regeneration within the time-frame of this event (Fig 1a).

The Western blots were performed with protein extracts from large tail sections and do not have the sensitivity to detect NF-*κ*B activity immediately adjacent to the amputation plane. The direct NF-*κ*B target and marker of inflammation ***Lim et al.*** (***2001***), *cox2*, is expressed along the amputation plane at 6 hpa and along the edge of tadpole tail regenerates throughout regeneration (Fig 1c). Expression of *cox2* was strong in stage 51 (regenerative) hindlimbs at 6 hpa and 1 day post-amputation (dpa) in the tissue beneath the wound epithelium, but was not observed at 2 dpa (Fig 1d). *Xenopus* limbs regenerate well early in development, but this capacity becomes lost as development progresses ***Dent*** (***1962***), allowing regenerative competency and incompetency to be studied in the same appendage (Fig 1e). Expression along the amputation plane at 1 dpa was reduced with decreasing regenerative capacity (Fig 1d, e). These data indicate a correlation between amputation plane expression of the NF-*κ*B target gene *cox2* in regenerating tails and limbs with eventual regenerative success.

### NF-*κ*B direct targets Nox2 and Nox4 are rapidly up-regulated in regenerating hindlimbs

NF-*κ*B also directly activates *nox1*, *nox2* and *nox4* ***Manea et al.*** (***2010***); ***Morgan and Liu*** (***2011***), which encode the catalytic subunit of different Nox complexes ***Miller et al.*** (***2006***). Since Nox is required for tail regeneration ***Love et al.*** (***2013***) and the NF-*κ*B target, *cox2*, is expressed in regenerating tails and hindlimbs, it was hypothesised that at least one of these NF-*κ*B-targeted *nox* genes would also be expressed during regeneration. Indeed, both *nox2* and *nox4* were expressed in regenerating stage 51 hindlimbs at all timepoints (Fig 2) spanning the timeframe of blastema formation ***Pearl et al.*** (***2008***). Expression of *nox2* was punctate and comparable to the previously described distribution of neutrophils and macrophages in regenerating hindlimbs ***Mescher et al.*** (***2013***), consistent with its expression being confined to these cells. Expression of *nox4* was observed around the injury and throughout the regenerate. No obvious expression of *nox1* was observed at any timepoints. These results suggest Nox2 and Nox4 as drivers of ROS production during regeneration. Preventing inflammatory cell recruitment to the injury does not significantly alter ROS production in regenerating tadpole tails ***Love et al.*** (***2013***), suggesting that Nox2 does not substantially contribute to overall ROS production during tail regeneration.

**Figure 2.**
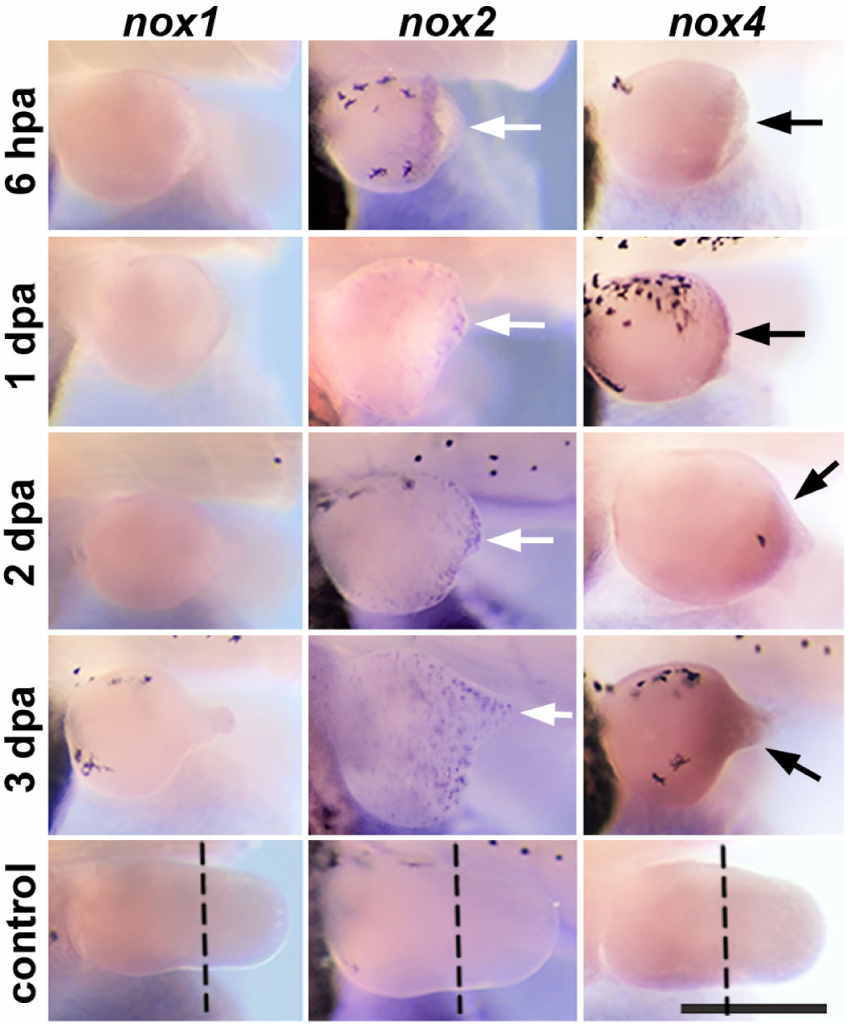
*nox2* and *nox4* are expressed in regenerating hindlimbs. *In situ* hybridisations, using indicated probes, of amputated stage 51 tadpole hindlimbs. Fixation times (hpa/dpa) are labelled. White arrows indicate punctate staining, black arrows indicated continuous staining and dashed lines indicate the amputation planes in unamputated controls. Scale bar: 500 µm.

### Resident microbes may activate regeneration of tadpole tails

*Xenopus laevis* tadpoles can regenerate their tails up to metamorphosis, but temporarily lose this ability during the ‘refractory period’ from stages 45 to 47 ***Beck et al.*** (***2003***). As ROS are required for successful tadpole tail regeneration, it was hypothesised that this refractory period could result from impaired ROS signalling. To see if acute exposure to exogenous ROS during the refractory period improved regeneration, tadpole tails were dipped in different concentrations of hydrogen peroxide (H_2_O_2_) after amputation at stage 46. Unexpectedly, the H_2_O_2_ dip reduced, rather than enhanced, regeneration at concentrations of 3% and 0.3% (Fig.3a). Interestingly, the control tadpoles underwent equally good regeneration, which is not expected in the refractory period. This prompted a re-validation of the refractory period under current culture protocols. A clear refractory period was not consistently observed in all tadpole cohorts (data not shown).

**Figure 3.**
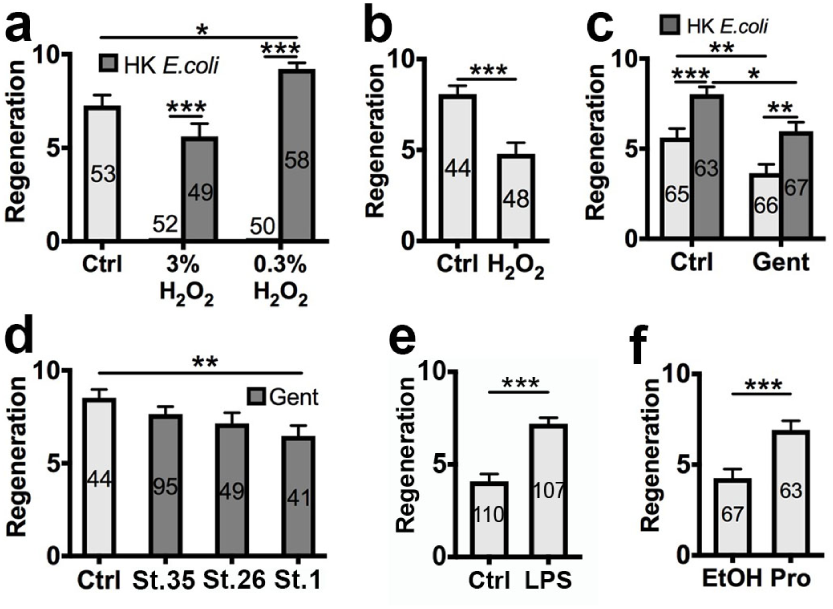
Regeneration is activated by bacterial ligands. Mean regeneration scores + s.e.m. of stage 46 tadpoles: **a**, with amputated tails dipped in the indicated concentration of H_2_O_2_ ± HK *E. coli* and untreated control; **b**, immersed in 0.3% H_2_O_2_ prior to tail amputation and untreated control; **c**, raised with or without gentamicin prior to tail amputation ± HK *E. coli*; **d**, raised with gentamicin beginning at the indicated developmental stage before tail amputation + HK *E. coli*; **e**, with tails amputated ± LPS; **f**, with tails amputated + prostratin or ethanol control. Sample numbers are indicated inside bars. *p* values are based on ordinal logistic regressions: * *p* < 0.05, ** *p* < 0.01, *** *p* < 0.001. Data for graphs can be found in supplemental file S1.

**Figure 4.**
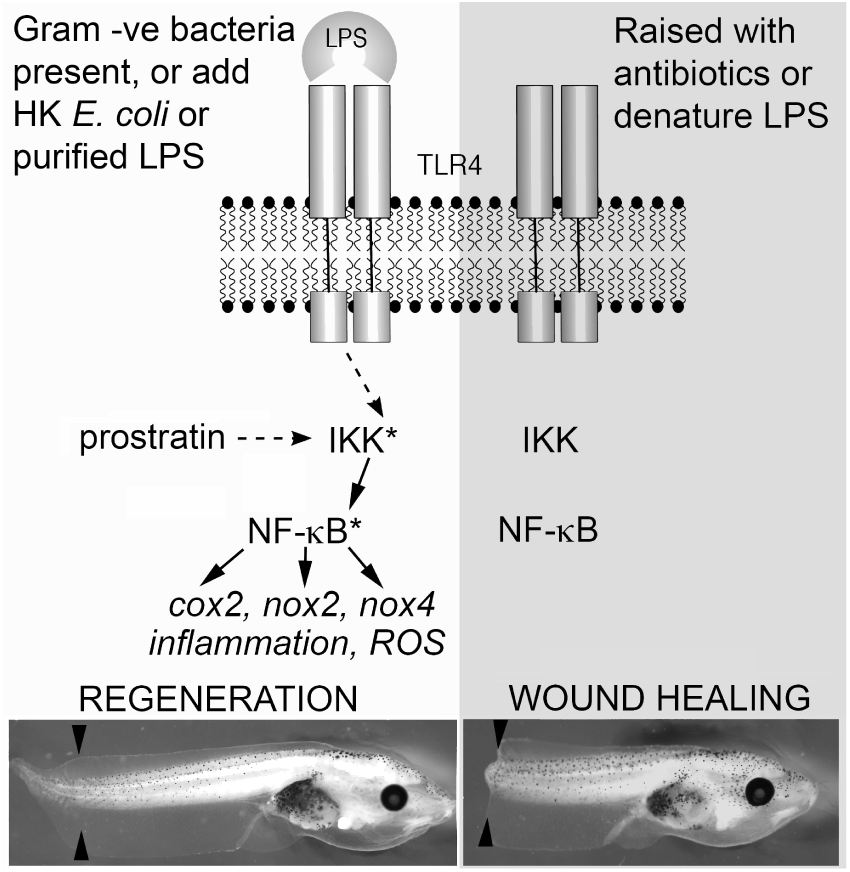
Proposed mechanism for how bacteria influence the outcome of regeneration following partial tadpole tail amputation. On the left side, regeneration is favoured when native microbiota are present (on the tadpole’s skin or in the media) or if heat killed *E. coli* or purified LPS are added to the media. The small molecule prostratin can mimic this by indirectly activating IKK. Downstream targets of NF-*κ*B could act to activate an inflammatory response and propagate the ROS signal reported by ***Love et al.*** (***2013***). On the right side, raising tadpoles in the antibiotic gentamicin, or treating tadpoles with H_2_O_2_ to denature LPS arising from native microbiota leads to a wound healing response. Asterisk denotes activated protein. Black arrowheads indicate the site of amputation at stage 46, representative tadpoles are shown 5 days after amputation.

The unexpected reduction in regeneration following H_2_O_2_ treatment prompted a search for an explanation. The NF-*κ*B activator, IKK, can be activated by the binding of ligands from microorganisms to toll-like receptors (TLR) ***Chow et al.*** (***1999***). Lipopolysaccharides (LPS) are present on the surface of Gram-negative bacteria and bind to TLR4, eliciting activation of NF-*κ*B via IKK ***Chow et al.*** (***1999***). LPS which have been treated with H_2_O_2_ no longer function as TLR4 ligands ***Cherkin*** (***1975***). It was hypothesised that dipping tails in H_2_O_2_ was inhibiting regeneration by disabling TLR ligands of bacteria around the wound, preventing TLR activation. In the absence of TLR activation, IKK may not be activated on amputation, potentially preventing NF-*κ*B activity and the ROS burst being sustained.

To test this, heat-killed (HK) *Escherichia coli* (Gram-negative bacteria with surface LPS ***Pradhan et al.*** (***2012***)) were added following tail amputation and H_2_O_2_ treatment (Fig 3a). Regeneration was greatly improved following addition of HK *E. coli*. Strikingly, tadpoles treated with 0.3% H_2_O_2_ and HK *E. coli* had significantly better regeneration than the untreated controls. To see if H_2_O_2_ treatment of tadpole skin, and not the wound itself, was sufficient to inhibit regeneration, stage 46 tadpoles were brie2y immersed in H_2_O_2_ prior to amputation (Fig 3b). Regeneration was significantly reduced with H_2_O_2_ pre-treatment, consistent with the hypothesis that naturally occurring bacterial ligands on the skin surface have an important role in initiating regeneration. The reduction in regeneration with H_2_O_2_ treatment varied between batches of tadpoles (data not shown), possibly due to varying bacterial loads.

To further test this hypothesis, tadpoles were raised with the broad spectrum antibiotic, gentamicin, to reduce the bacterial load on the skin (Fig 3c). Tadpoles raised with gentamicin had low regeneration scores after amputation at stage 46, which were significantly increased by addition of HK *E. coli*. Interestingly, with added HK *E. coli*, the gentamicin treated tadpoles had significantly lower regeneration scores than their siblings raised without gentamicin. Gentamicin reduced the regenerative potential of tadpole tails, even with addition of HK *E. coli*, and this reduction was greater the earlier gentamicin treatment began (Fig 3d). Purified LPS also significantly improved regeneration of stage 46 tadpole tails (Fig 3e), as did the protein kinase C activator, prostratin (Fig 3f), which activates IKK intracellularly, independent of TLRs ***Williams et al.*** (***2004***). These results reveal the involvement of ligands binding to TLRs in the activation of regeneration, possibly through the IKK pathway.

## Discussion

Regeneration of the *Xenopus* tadpole tail can be divided into three distinct phases: an early wound healing stage, which takes place from 0- 6h, an intermediate phase in which the regeneration bud is formed (around 24h) and a late phase, from 48h to 1 week, in which replacement of lost tissues is completed ***Love et al.*** (***2013***); ***Beck et al.*** (***2009***); ***Ferreira et al.*** (***2016***). The importance of ROS, particularly H_2_O_2_, is increasingly being recognised as a critical factor influencing regenerative success ***Chen et al.*** (***2014***). The production of ROS is the earliest known response to amputation, with detection possible within a few minutes and a gradient rapidly established which is thought to attract immune cells such as macrophages to the wound site ***Love et al.*** (***2013***). Here, we show that NF-*κ*B, a key regulator of the immune response, is rapidly translocated to the nucleus following tail amputation, and that the direct target *cox2* is also rapidly expressed at the wound surface. The apparent rapid activation of NF-*κ*B not only provides a biochemical link between ROS production and the activation of the immune response, but also provides a possible mechanism for regulating continuous ROS production via a feedback loop. NF-*κ*B targets include the NADPH oxidase genes and we show here that *nox2* (in macrophages) and *nox4* expression (in distal cells) are also rapidly upregulated. NF-*κ*B may therefore play a role in modulating ROS levels, which others have shown to be critical for the active re-polarisation of bioelectric circuitry that contributes to regenerative outcomes ***Ferreira et al.*** (***2016***).

While others have shown that physiological levels of exogenous H_2_O_2_ applied in the first 24 hours (but not if left on for the whole process) can improve refractory regeneration ***Ferreira et al.*** (***2016***), here we show that brief exposure to higher levels of exogenous H_2_O_2_ has the opposite effect, and that this can be rescued by heat killed Gram-negative bacteria (*E. coli*) or purified LPS, suggesting a role for Gram-negative tadpole skin microbiota in determining the outcome of tail regeneration in tadpoles from stage 45. LPS binds to TLRs, which can in turn activate NF-*κ*B, but treatment with H_2_O_2_ prevents this binding ***Cherkin*** (***1975***). Gentamicin is a broad spectrum bacteriostatic antibiotic commonly used by researchers raising *Xenopus* embryos to reduce early deaths. The refractory period may therefore be revealed in a sterile laboratory environment. The refractory period ends at stage 48 which can only be reached in fed tadpoles, which are no longer in a sterile environment and so could regain regenerative capacity through TLR pathway activation.

TLR4 is limited to only a few cell types, including macrophages ***Waltenbaugh et al.*** (***2008***), so TLR4 would not be sufficient to activate NF-*κ*B-mediated regeneration in other cell types. TLR4 pathway activation in macrophages leads to the production and release of pro-inflammatory cytokines including TNF and IL-1 ***Zheng et al.*** (***2012***) which have receptors on most cell types ***Rothwell et al.*** (***1997***); ***Dinarello*** (***1998***); ***Galeone et al.*** (***2013***). Binding of these cytokines to their respective cell-surface receptors leads to the activation of IKK ***Hayden and Ghosh*** (***2004***) and could consequently spread the activation signal to many cell types. TNF is important for skeletal muscle regeneration ***Li and Schwartz*** (***2001***); ***Warren et al.*** (***2002***) and liver regeneration ***Diehl et al.*** (***1994***, 1995) and essential for zebrafish fin regeneration ***Nguyen-Chi et al.*** (***2017***). The recruitment of macrophages may thereby be important for TLR-dependent regeneration. Interestingly, macrophage removal prevents regeneration of salamander limbs ***Godwin et al.*** (***2013***). The regenerative competency of pre-refractory tadpole tails, raised in a sterile environment ***Beck et al.*** (***2003***), indicates that they must have an alternative, TLR-independent, and perhaps also IKK-independent, mechanism of regeneration. Reducing inflammatory cell migration does not prevent tail regeneration in stage 43 (pre-refractory-equivalent) *Xenopus tropicalis* tadpoles ***Love et al.*** (***2013***).

This study suggests a role for NF-*κ*B in regeneration and a mechanism for its continual activity in blastemal cells through a positive-feedback loop involving Nox4 and ROS. A method to initiate this positive-feedback loop in human cells may have the potential to improve our regenerative capabilities.

## Methods and Materials

### Animal ethics

All animal experiments were approved by the University of Otago Animal Ethics Committee under protocol AEC56/12.

### Tail regeneration

Tadpoles were raised in petri dishes (40-60 per dish) in 0.1× Mark’s modified Ringer’s (MMR) at 24°C. They were divided evenly between treatment groups either before (when using pre-amputation treatment) or after amputation. A sharp scalpel blade was used to amputate 30% of the tail under MS222 (1/4000 w/v) anaesthesia (>stage 44) or no anaesthesia (<stage 45). Following any post-amputation treatment, each group of tadpoles was divided into at least three petri dishes. Tadpoles were kept at 24°C for 7 days with daily feedings of spirulina and 50% 0.1x MMR changes. Regeneration was scored as follows: 0=no regeneration, 5=partial regeneration and 10=complete regeneration.

### Hindlimb regeneration

Stage 51-55 tadpoles were anaesthetised with MS222 (Sigma, 1/4000 w/v) and placed on a moistened paper towel. One hindlimb was amputated using Vannas iridectomy scissors at the approximate level of the knee. Tadpoles were transferred to dechlorinated water with air bubbled through an aeration stone. Tadpoles were kept in the aquarium at 25°C for the stated time (or for 1 month when regeneration was scored). Regeneration scores were assigned as follows: 0=stump, 0.5=spike, 1-5=the number of regenerated digits.

### Heat killed *E. coli*

TOP10 *Escherichia coli* were grown in 10ml of Luria broth to stationary phase before centrifugation at 13,000 rpm and resuspension in 1x PBS three times. They were then heated to 70°C for 1 h, centrifuged and resuspended in 2 mL of 0.1x MMR. As indicated, the solution was added at a dilution of 1:100 in 0.1x MMR for 1 h following amputation.

### H_2_O_2_ dip

Stage 46 tadpoles were anaesthetised with MS222 and tails were amputated using a sharp scalpel blade. Tadpoles were placed in 0.1× MMR (MS222 and H_2_O_2_ form a toxic product, so tadpoles must be removed from MS222 before H_2_O_2_ dip) and individually sucked headfirst into an unmodified 3 mL plastic pipette with tails projecting out the end. Tails were dipped in a H_2_O_2_ solution for 3 s and the tadpoles were placed in fresh 0.1× MMR three times before being transferred to their post-H_2_O_2_ treatments.

### H_2_O_2_ immersion

Stage 46 tadpoles were immersed in 0.3% H_2_O_2_ in 0.1× MMR for 2 m and transferred to fresh 0.1× MMR and anesthetised with MS222 prior to tail amputation.

### DPI treatment

Stage 43 tadpoles were treated with 2 µM diphenyleneiodonium chloride (DPI; Sigma; 2 mM in DMSO stock) or 0.1% DMSO (in 0.1× MMR) for 1 h before and 1, 3, 6 or 12 h after tail amputation.

### LPS and prostratin treatments

Stage 46 tadpoles (raised at a low density of approx. 40 per dish) were anaesthetised with MS222 and tails were amputated using a sharp scalpel blade. Tadpoles were treated with 50 µg.mL^−1^ lipopolysaccharides (LPS) from *E coli* 0111:B4 purified by phenol extraction (Sigma; 5 mg.L^−1^ in water stock) with untreated control for 1 h, or 10 µM prostratin (Sigma; 4 mg.mL^−1^ in ethanol stock) with 0.1% ethanol control, for 40 m.

### Gentamicin treatment

In summary, tadpoles used for gentamicin experiments were raised at 17°C and placed in petri dishes (40-60 per dish) containing 100 µg.L^−1^ gentamicin in 0.1× MMR from the stage indicated, or if no stage is indicated from just after the embryos were sorted at stage 2-6. Embryos were moved to new petri dishes containing fresh gentamicin every two days. Following amputation, tadpoles were incubated at 24°C and remained in gentamicin for a further day.

### Cytoplasmic and nuclear protein extractions and Western blots

Stage 55 tadpole tails (four per timepoint) were amputated (approx. 40% of tail length) with a sharp scalpel blade under MS222 anaesthesia. After 0, 1, 3 or 6 h, tadpoles were re-anaesthetised and a further 3 mm (20% of the remaining tail) was amputated. For DPI treatment, tadpoles were incubated with 2 µM DP1 (0.1% DMSO) for 1 h prior to the first amputation until the second amputation. The four 3 mm sections for each timepoint and treatment were homogenised in a bead beater with CER I (NE-PER) reagent, Halt protease inhibitor cocktail (Thermo Fisher) and EDTA. Cytoplasmic and nuclear protein fractions were extracted using NE-PER Nuclear and Cytoplasmic Extraction Reagents (Thermo Fisher) following the manufacturer’s protocol. Nuclear extracts were diluted 1:1 with PBS. Samples were prepared with sample loading buffer and heated to 100°C for 3 m. The samples (10 µL) were run on 12.5% SDS-PAGE gels along with a Precision Plus Protein dual colour standard (Bio-Rad) and transferred to PVDF membranes. The membranes were blocked with 2% BSA in PBS/tween (25 m) and incubated with 1:200 rabbit-anti NF-B p65 polyclonal antibody (Pierce) (2 h) and 1:5000 Goat anti-rabbit IgG (H+L)-HRP conjugate (Bio-Rad) (1 h). Reactivity was detected with Clarity Western ECL Substrate (Bio-Rad).

### Whole mount *in situ* hybridisation

*Coding regions of cox2 (primers: 5’ gctctagaatgatcgtactacccgc, 3’ cggtaccttaaagttcggatgtgtgc)*, nox1 (primers: 5’ tcctgttttccagggcagtg, 3’ agattggacgcccatagctg), *nox2* (primers: 5’ ctatgacgagggcgaagcat, 3’ tcatcccagccagtgaggta) and *nox4* (primers: 5’ taggcaggaatccagtgatgg, 3’ cactcccgcaacagaagtga) were amplified from stage 12 *X. laevis* embryo cDNA using Pwo DNA Polymerase (Roche) and ligated into the TOPO Cloning site of the Pcr4-TOPO plasmid (Invitrogen Life Technologies). Insertions were verified by DNA sequencing. Plasmids were linearised with NotI and digoxigenin-labelled RNA probes were transcribed using T3 polymerase with a DIG-NTP mix (Roche). Templates were removed using DNase I (Roche) and the probes were precipitated with 2.5 M LiCl. Whole-mount in situ hybridisations were performed as previously described in Pearl et al (12).

## Acknowledgments

We would like to thank Adam Middleton and Robert Day for comments on the manuscript and Esther Pearl for prior cloning of *X.laevis Cox2*. TFB was supported by a University of Otago PhD studentship.

